# Plasma-derived cell-free DNA methylomes might not enable detection and discrimination of intracranial tumors

**DOI:** 10.1101/2021.06.02.446405

**Authors:** Kristoffer Vitting-Seerup

**Affiliations:** The Bioinformatics Section, Department of Health Technology, The Technical University of Denmark (DTU), Denmark

## Abstract

Intracranial tumors are both hard to detect and diagnose, resulting in poor patient outcomes. For that reason, non-invasive methods enabling detection and discrimination of intracranial tumors have significant clinical potential. Recently Nassiri et al. propose plasma-derived cell-free DNA methylomes as such a method. Here I show that the results have been misinterpreted, and for many comparisons, no evidence supporting the conclusions are actually presented. While my analysis highlights the potential of plasma-derived cell-free DNA methylomes, the evidence provided by Nassiri et al. is simply currently insufficient.

## Main Text

Intracranial tumors are both hard to detect and diagnose, resulting in poor patient outcomes. For that reason, non-invasive methods enabling detection and discrimination have significant clinical potential. Recently Nassiri *et al*. proposed plasma-derived cell-free DNA methylomes as such a method^1^. Here I show that Nassiri and coauthors have misinterpreted their results and currently provide little or no evidence supporting the conclusion that plasma-derived cell-free DNA methylomes enable detection and discrimination of intracranial tumors.

Intracranial tumors are notoriously difficult to detect since most symptoms, such as headaches, are nonspecific and frequently occur in healthy people^2^. Therefore, these tumors often go unnoticed, resulting in late discovery and worse patient outcomes^2^. When intracranial tumors are discovered, highly invasive intracranial surgery is needed to confirm and elaborate on the diagnosis^1^. Non-invasive approaches capable of detecting and discriminating intracranial tumors are central to both screening and diagnosis efforts and could ultimately improve patient outcomes.

Nassiri *et al*. recently reported they could detect and discriminate intracranial tumors using plasma-derived cell-free DNA methylomes^1^. Their study created plasma-derived methylome profiles for 220 patients with various types of intracranial tumors. These methylome profiles were then combined with previously published methylome profiles (n=447) from both healthy controls and other cancer types. Nassiri *et al*. used this dataset to analyze whether: 1) gliomas can be distinguished from extracranial samples; 2) whether intracranial tumor types can be distinguished from each other. In both cases, they frame the analysis as a one-vs-rest classification problem (e.g., glioma vs. all extracranial samples) and train random forest models to solve the problem^1^. For both comparisons, they use nested cross-validation and only report performance via Receiver Operator Curve (ROC) summarized as Area Under receiver operator Curve (AUC)^1^.

Unfortunately, the combination of an imbalanced dataset (e.g., 60 gliomas vs. 447 extracranial samples) and only reporting performance with ROC/AUC can be misleading. While ROC and AUC are undoubtedly among the most reliable performance estimators^3–6^, it is widely recognized that ROC/AUC is inadequate for imbalanced datasets^3,5,7–9^. ROC analysis simply provides an over-optimistic evaluation for imbalanced datasets that gets worse the more imbalanced the dataset^3^. For a detailed discussion of this phenomenon, please refer to, e.g., Saito *et al*.^8^ or Nature’s “Points of Significance” article about classification evaluation^6^. The widespread acceptance of this problem is reflected in many approaches developed to handle the issue. Among the solutions applicable before/during model training are down-sampling or re-sampling of datasets to create equal-sized classes or adding increased weights to the minor class to ensure the model pays more attention to the minor class^7^. In addition, after a model has been trained on an imbalanced dataset, it should not only be evaluated using sensitivity/specificity (jointly creating ROC) but also via performance measures such as precision, False Discovery Rate (FDR), or Precision-Recall Curves (PRC)^8,10,11^. Due to the complementarity of the two approaches, state-of-the-art is currently to evaluate a model using both ROC and PRC curves^6^.

In the context of the Nassiri *et al*. paper, the above-described shortcoming of ROC means that many glioma samples could mistakenly be predicted as “non-glioma” without noticeably affecting the AUCs. In this paper, I investigated if this was the case. Specifically, I reanalyzed the actual predictions made by Nassiri *et al*. (available in their supplementary material) using additional performance metrics. Unfortunately, I find that the results supporting the main conclusions in Nassiri *et al*. are either missing or flawed.

The problem with imbalanced datasets is most prominent in the analysis of intracranial samples. Here Nassiri *et al*. analyze 161 intracranial cancer samples originating from 6 different cancer types: 41 “IDH mutant gliomas”, 22 “IDH wild-type gliomas”, 60 meningiomas, 9 hemangiopericytomas, 14 “low-grade glial–neuronal tumors”, and 15 “brain metastases”. To determine if one cancer type could be distinguished from the other cancer types, they train a model to distinguish, e.g., the 14 low-grade glial–neuronal tumors from the 147 other cancer samples (8.7% vs. 91.3%). As reported in Nassiri *et al*., the average resulting AUC for this comparison is 0.93, indicating almost perfect performance. However, when I inspected the results provided in the supplementary material more thoroughly, I found the high AUCs originate from classifying all samples as ‘other’. In other words, no samples were classified as low-grade glial–neuronal tumors (Figure 1, leftmost plot), meaning the results are clinically unusable. I find this exact pattern for four of the six cancer types (Figure 1), with the other cancer types not being much better. In other words, for most cancer types, the authors literally provide no support for the hypothesis. Hence Nassiri *et al*.^1^ currently do not provide the evidence to conclude that plasma-derived cell-free DNA methylomes enable discrimination of intracranial tumors.

**Figure 1:**
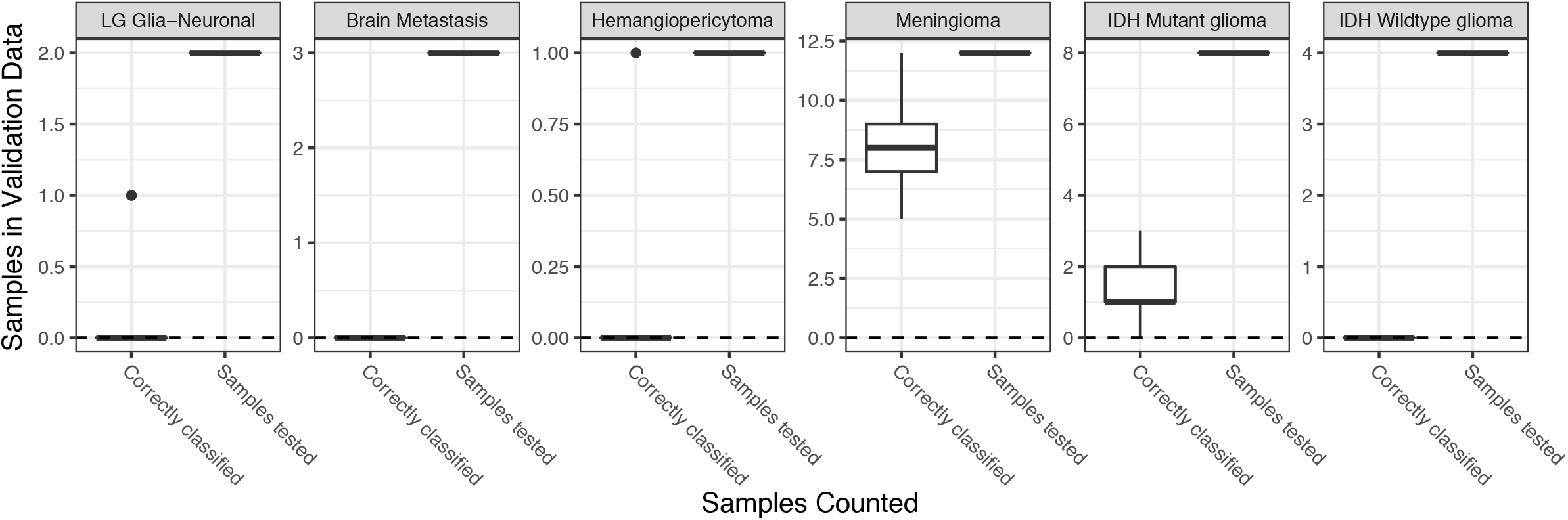
Distinguishing intracranial tumors. For each tumor type (sub-plots), the number of samples (y-axis) tested and the number of samples correctly classified (x-axis) for the 50 iterations of prediction on held-out data provided by Nassiri *et al*.

Apart from the intracranial analysis, Nassiri *et al*.^1^ also seek to determine if intracranial tumors can be distinguished from extracranial samples. Here a one-vs-rest approach is used to compare glioma samples to a dataset containing a range of other cancer types as well as healthy controls. This corresponds to comparing 60 gliomas samples (11.8%) to 447 non-gliomas samples (88.2%), meaning it is also an imbalanced dataset. As reported in Nassiri *et al*., this comparison yields an average AUROC is 0.99, which considering an AUROC score of 1 means perfect classification, seems very impressive. Such a conclusion would, however, be a misinterpretation of the AUCs. When I analyzed the results provided in the supplementary material with additional metrics, I found the predictions to have a False Discovery Rate of 18.2% (Figure 2A). This means that approximately 1 in 5 glioma patients would not be identified by the models proposed by Nassiri *et al*. (Figure 2A).

**Figure 2:**
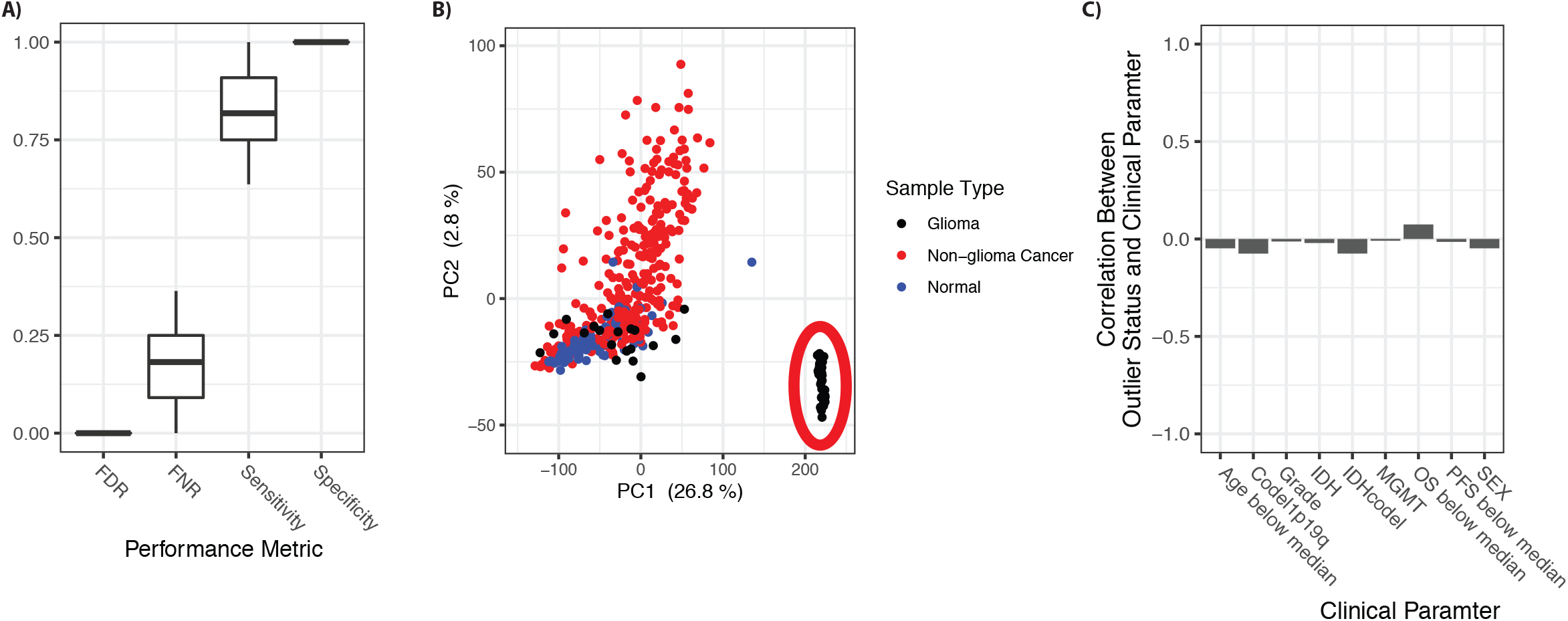
Detecting intracranial tumors. **A)** The x-axis indicates the different performance metrics calculated directly from the predictions provided in Nassiri et al. Y-axis indicates the performance for the 50 iterations of prediction on held-out data provided by Nassiri *et al*.. **B)** PCA plots of the 25,000 most informative methylated regions show the clustering of glioma and extracranial samples. Samples are colored by type, and a red ellipse highlights the glioma outliers. **C)** Correlation between glioma outlier status (as defined in B) and clinical metadata. OS: Overall Survival. PFS: Progression Free Survival.

There is, however, another problem with the analysis of gliomas vs. extracranial samples. For this analysis, Nassiri *et al*.^1^ combine three distinct datasets originating from different laboratories (and distinct points in time). When comparing different datasets, they almost always result in batch effects where systematic variation between samples is introduced because of differences in the laboratory conditions (e.g., temperature), personnel, etc.^12^. We know batch effects exist in the data analyzed by Nassiri *et al*. as they point to batch effects as the reason the data is analyzed as two individual cohorts instead of one combined^1^. It is, however, currently unclear if or how Nassiri *et al*. have corrected other potential batch effects as this is not described in the method section nor the supplementary code. It does, however, appear that batch effects have not been adequately handled as the glioma samples appear as two distinct clusters in a PCA analysis (Figure 2B) (also seen in Figure 1G in Nassiri *et al*.^1^). This clustering could result from a batch effect - a hypothesis supported by the clusters not being explained by the available clinical meta-data (Figure 2C). Unless an unpublished clinically relevant feature explains this outlier group, it should most likely be removed using bioinformatic tools such as sva^13^. The combination of the sub-standard performance (Figure 2A) with the uncertainty of the reliability due to batch effects (Figure 2B-C) means that Nassiri *et al*.^1^ currently do not provide the evidence to conclude that plasma-derived cell-free DNA methylomes enable detection of intracranial tumors.

In summary, while the data analyzed by Nassiri *et al*.^1^ might have the potential, the current analysis simply provides insufficient evidence to conclude that plasma-derived cell-free DNA methylomes enable detection and discrimination of intracranial tumors.

## Methods

The processed data, the scripts created, and the binary classification of samples tested in Nassiri *et al*. were downloaded from their Zenodo repository (www.doi.org/10.5281/zenodo.3715312). A Rmarkdown document reproducing the analysis presented here can be found on Figshare (http://doi.org/10.6084/m9.figshare.14406866.v1). Performance metrics were calculated as defined in Saito *et al*.^3^ directly from the binary classification provided in Nassiri *et al*. Zenodo repository (see Rmarkdown).

